# HieVi: Protein Large Language Model for proteome-based phage clustering

**DOI:** 10.1101/2024.12.17.627486

**Authors:** S. Panigrahi, M. Ansaldi, N. Ginet

**Affiliations:** Phage cycle and bacterial metabolism – Laboratoire de Chimie Bactérienne – UMR7283 CNRS/Aix-Marseille Université, Marseille, France

**Keywords:** Protein language model, Machine learning, Bacteriophages, Viral taxonomy, Comparative genomics

## Abstract

Viral taxonomy is a challenging task due to the propensity of viruses for recombination. Recent updates from the ICTV and advancements in proteome-based clustering tools highlight the need for a unified framework to organize bacteriophages (phages) across multiscale taxonomic ranks, extending beyond genome-based clustering. Meanwhile, self-supervised large language models, trained on amino acid sequences, have proven effective in capturing the structural, functional, and evolutionary properties of proteins. Building on these advancements, we introduce HieVi, which uses embeddings from a protein language model to define a vector representation of phages and generate a hierarchical tree of phages. Using the INPHARED dataset of 24,362 complete and annotated viral genomes, we show that in HieVi, a multi-scale taxonomic ranking emerges that aligns well with current ICTV taxonomy. We propose that this method, unique in its integration of protein language models for viral taxonomy, can encode phylogenetic relationships, at least up to the family level. It therefore offers a valuable tool for biologists to discover and define new phage families while unraveling novel evolutionary connections.

## Introduction

Self-supervised Large Language Models (LLM) based on transformer architecture have been successful in encoding contextual information content of a word (or more generally a token) in a sentence. Various methods have been developed using the learnt vector representations of the tokens (or embeddings) for classification of sentences and further for clustering of texts^1^. Similarly, protein Language Models (pLMs) treat amino-acid sequences as tokens and, due to availability of large datasets of proteins, have been trained in a self-supervised manner to learn contextual information and evolutionary fitness of amino-acids in a sequence. This information is encoded in the embeddings, which are vectorial representations of this information in an abstract continuous space. The embeddings and attention maps from different layers of the model have been shown to also encode structural information of a protein^2–4^. There are now few such pre-trained pLMs available to the public, for example Evolutionary Scale Modeling 2 (ESM-2)^5^ or ProtT5-XL-U50^6^. These models are trained on the large protein UniRef50 dataset^7^, thus capturing the evolutionary fitness of proteins in the embeddings. ESM shows that protein representations extracted from the pre-trained pLM cluster into groups with similar structure and align with previously annotated protein families^3^. Thus, proteins which are closer in the protein representations space are likely to be functionally similar.

We posit that protein representations generated from pLMs can be extended to entire proteomes to compare and classify biological entities. By proteome we mean the set of distinct proteins that is encoded by any single biological entity’s genome (*e.g.*, a eukaryotic cell, an archaeon, a bacterium or a virus). One can view a proteome as the summation of functions acquired through evolution and one way to apprehend evolutionary relationships between biological entities can be done by comparing their proteomes.

Here we focused on a particular class of biological entities, the viruses of bacteria (bacteriophages or phages), that are defined by rather small proteomes compared to bacterial, archaeal and eukaryotic ones. Phages are diverse in terms of nucleic acid supporting the genetic information (*e.g.*, DNA, RNA, single-stranded or double-stranded), their gene content as well as the range of bacterial hosts they infect. Phages have been extensively studied for more than a century and there exists an abundant scientific literature covering all aspects of phage studies. The INPHARED database (September 2024 release) contains 29,759 reportedly complete, annotated prokaryotic virus genomes^8^. This number is increased by a factor 30 with metagenomic data in the PhageScope database^9^. Comparing and classifying such a high and exponentially increasing number of genomes is a key issue in the field. Phage taxonomy is a challenging task owing to the propensity of phages for recombination between viruses or with their hosts. Phage genome organization is described as mosaïc, with sets of genes regrouped in modules related to certain functions such as capsid or tail assembly, genome replication or cell lysis^10–12^. Such modules can be swapped between phages through recombination, rendering phage evolutionary history particularly challenging. Typical examples are lambdoid phages exemplified by the famous *Escherichia coli* phage Lambda. Furthermore, in contrast with other biological entities, there aren’t any universally shared genetic traits among viruses such as the 16S rRNA gene in bacteria. Viral taxonomy has to rely on multiple tools with different results as illustrated by Barylski et *al.* study of the phage *Herelleviridae* family and manual curation of taxonomy experts^13^.

The current viral classification including bacteriophages is kept separated from other biological entities and maintained by the ICTV^14^. The long history of phage classification reflects progresses in phage biology and investigation techniques over a century. This classification is currently undergoing a profound overhaul enabled by whole genome and proteome pipeline analyses with on-going efforts towards a unified taxonomy^15^. State-of-the art viral comparative genomics relies now on whole genome and proteome approaches such as vConTACT v2.0, VipTree or GRAVITy^16–18^. The ICTV classification has undergone a profound reshaping in 2022 enabled by the above-mentioned approaches^19^. Whereas species and genera are now clearly defined on Average Nucleotide Identity (ANI) thresholds (95% for species and 70% for genus demarcations), whole genome DNA comparisons usually fail to capture evolutionary relationships to define higher taxonomic ranks. Subfamily and family demarcation criteria are still non-homogenous as they depend on available data, bioinformatic analyses available at the time of their definition and their various combinations by different authors. As an example the *Autographivirinae* subfamily - promoted at the family level in 2021 as *Autographiviridae* - has been proposed in 2009 on the following criteria : “*encode their own RNA polymerase and share a common general genome organization, with genes encoded solely on the Watson strand*” (ICTV proposal 2008.020-023B.v3). This definition is biased towards a single, specific biological activity. The *Ackermannviridae* family and its internal structure relies on DNA sequence identity, shared prokaryotic virus orthologous groups (pVOG) and single protein phylogenies (terminase large subunit and tail-sheath proteins) studies. To the best of our knowledge, the most comprehensive study for phage family definition is exemplified in the above-mentioned work of Barylski et *al.*^13^ Using a double-stranded DNA (dsDNA) similarity network generated with vConTACT v2.0, the authors proposed a new taxon for a group of 93 dsDNA tailed phages, the *Herelleviridae* family. By combining genome-based and proteome-based analyses, gene synteny analysis and marker proteins phylogenies, the authors could refine the internal structure of this new taxon into five defined subfamilies and 15 genera. This work is another illustration of the on-going effort to clarify taxon demarcations in phage taxonomy using genome-wide approaches.

The revolution enabled by massive parallel DNA sequencing during the past ten years and the ensuing development of metagenomics resulted in an exponentially increasing number of available viral sequences. As an illustration, the online bacteriophage database PhageScope registers 873,718 bacteriophage genomes and 43,088,582 annotated proteins (November 2024 release)^9^. These numbers cannot but increase exponentially as new environments are being intensively sampled and their viromes sequenced. Hence, there is a crucial need for new comparative genomics approaches to grasp the prokaryotic virosphere in a more global, systematic and scalable way.

We present here an approach we named HieVi (for Hierarchical Viruses) using pLM-based whole proteome clustering to address the challenges posed in viral comparative genomics and classification. Since phage comparative genomics relies now on shared protein functions, we hypothesized that a simple average over all protein representations from a phage proteome encodes the information about distribution of protein functions within that proteome. HieVi encodes each phage proteome by a single vector, a Mean Phage Representation (MPR). We generated MPRs for the 24,362 complete and annotated prokaryotic viral genomes present in the HieVi and clustered them using a density-based clustering algorithm. This allowed us to construct a HieVi hierarchical tree computed from the pairwise Euclidean distance matrix between MPRs. We observed that the HieVi hierarchical tree topology contained multi-scale taxonomic information. From the largest perspective to the finer grain, we observed that viruses first cluster in the tree according to the realm and kingdom they belong to, then that phages belonging to the same ICTV family are, in many cases, clustered in the same branch and finally that phages generally also cluster within this branch according their ICTV genus and subfamily annotations. We showed that HieVi topology is in good accordance with established phage phylogenies at multiple scales. We documented these observations with a few known phage families and provided an example of how HieVi can be used to refine current taxonomy and correct inappropriate assignments. We finally propose that HieVi can help define new phage taxa and discover unsuspected evolutionary relationships. In a broader perspective, this study is the first example of protein language model implementation to compare biological entities. HieVi can be extended to more complex organisms such as bacteria, archaea or eukaryotes, computational resources, biological diversity and expert validation being the only limits for this.

## Methods

### Viral proteome dataset

We used the September 2024 release of the INPHARED prokaryotic viral database maintained and regularly updated by Millard’s lab^8^. This dataset comprises 29,759 annotated genomes, mostly from the RefSeq and Genbank databases. We filtered this database for unique accessions, resulting in the HieVi dataset with 24,362 genomes and 2,166,614 proteins. The authors regularly curate their database from incomplete genomes. To ensure the best possible consistency across the dataset, they also performed gene calling on their entire database with the same gene caller. This dataset includes mostly phages but also viruses infecting archaea and unclassified hosts. It also covers a wide range of proteome sizes (Table 1).

**Table 1:** HieVi dataset. A) Number of viruses sorted by their prokaryotic host super-kingdom. B) Proteome size distribution (number of CDS per genome). Total number of genomes *n* = 24,362.

| (A) Prokaryotic host |  |  | (B) Proteome size |  |  |
| --- | --- | --- | --- | --- | --- |
| Bacteria | Archaea | Unclassified | < 10 | 50 - 100 | > 200 |
| 20,995<br>(86.2%) | 295<br>(1.2%) | 3,075<br>(12.6%) | 14.2% | 47.7% | 13.0% |

This dataset covers all taxonomic ranks present in the current ICTV prokaryotic viral taxonomy (Table 2).

**Table 2:** Taxonomic description of the HieVi dataset. Taxonomic ranks are defined by the ICTV and displayed in hierarchical order (the realm being the highest taxonomic rank). The four top annotated taxa were displayed for each taxonomic rank. ICTV accepted proposals defining current viral taxa are available through the ICTV website (https://ictv.global/taxonomy).

|  | ICTV | Unclass. | Top 1 | Top 2 | Top 3 | Top 4 |
| --- | --- | --- | --- | --- | --- | --- |
| Realm | 6 | 4.9% | *Duplodnaviria*<br>79.4% | *Monodnaviria*<br>14.9% | *Varidnaviria*<br>0.5% | *Riboviria*<br>0.2% |
| Kingdom | 9 | 4.9% | *Heunggongvirae*<br>79.4% | *Sangervirae*<br>13.6% | *Loebvirae*<br>1.2% | *Bamfordvirae*<br>0.5% |
| Phylum | 9 | 4.9% | *Uroviricota*<br>79.4% | *Phixviricota*<br>13.6% | *Hofneiviricota*<br>1.2% | *Preplasmiviricota*<br>0.5% |
| Class | 9 | 4.9% | *Caudoviricetes*<br>79.4% | *Malgrandaviricete*<br>13.6% | *Faserviricetes*<br>1.2% | *Tectiliviricetes*<br>0.5% |
| Order | 15 | 83.8% | *Petitvirales*<br>13.6% | *Tubulavirales*<br>1.2% | *Kalamavirales*<br>0.4% | *Crassvirales*<br>0.3% |
| Family | 77 | 57.1% | *Microviridae*<br>13.6% | *Autographiviridae*<br>7.1% | *Straboviridae*<br>3.8% | *Herelleviridae*<br>2.3% |
| Subfamily | 108 | 67.5% | *Studiervirinae*<br>3.5% | *Tevenvirinae*<br>2.6% | *Bullavirinae*<br>2.2% | *Bclavirinae*<br>1.6% |
| Genus | 1388 | 34.5% | *Microvirus*<br>2.8% | *Gequatrovirus*<br>1.5% | *Tequatrovirus*<br>1.4% | *Fromanvirus*<br>1.4% |

Table 2 illustrates the current challenges in viral taxonomy with about ∼57% unclassified viruses at the family level and ∼34% at the genus level. This table also points at several sampling biases inherent to current phage genome databases. For instance, dsDNA tailed-phages (*Caudoviricetes* class) represent ∼79% of INPHARED genomes whereas few single-stranded RNA (ssRNA) phages (*Riboviria* realm) are recorded (0.2%). This is mostly due to historical reasons (*e.g.*, focus on human pathogenic bacteria), practical reasons (*e.g.*, usage of cultivable lab strains for phage isolation) or methodological reasons (*e.g.*, sequencing techniques favoring dsDNA viruses). These biases are being progressively reduced, thanks to metagenomic and metaviromic approaches on samples from extraordinarily diverse environments (*e.g.*, oceans, glaciers, clouds, sediments or microbiota).

#### Phage similarity network generated by vConTACT v2.0

We generated with the HieVi dataset a phage similarity network using state-of-the-art vConTACT v2.0 pipeline based on shared protein clusters^16^. Diamond was used for protein sequence all-versus-all alignments. Protein clusters (PCs) and Viral clusters (VCs) were generated by MCL and ClusterOne, respectively. All other parameters were set to default values. The 2,166,614 viral protein sequences included in our dataset were first grouped into 125,242 PCs within which proteins are expected to perform similar functions. Among the 24,362 viruses in the dataset, 20,784 were clustered into 3,209 VCs where viruses are grouped according to PCs they share. The remaining 3,578 genomes either overlap with two VCs (2,044) or were defined as singleton (311) and outliers (1,223) by the pipeline. vConTACT v2.0 results for each accession in the HieVi dataset are summarized in the Supplementary Excel file HieVi_annotation_Sept2024.xlsx.

#### Pre-trained protein Language Model ESM-2

ESM-2 (Evolutionary Scale Modeling 2) is a protein Language Model (pLM) that leverages transformer architectures to learn amino-acid embeddings capturing biochemical and functional information within a protein^5^. It has been shown that ESM- 2 learns protein representations that encode the evolutionary and structural properties of proteins without explicit structural data. The protein representations align with a protein family and function, making them useful for downstream functional classification tasks such as predicting sub-cellular localization and catalytic activity.

In this study, we used the ESM-2 pre-trained model (esm2_t36_3B_UR50D) which has been trained on the dense and highly diverse protein dataset Uniref50 (UR50, version 2021/04) available at the Uniprot database (https://www.uniprot.org/). This pre-trained model currently contains 3 billion parameters, has 36 layers with an embedding dimension of 2,560.

Our starting viral dataset consisted in 2,166,614 protein sequences and necessitated a GPU with sufficient memory capacity to load the model and perform inference efficiently. For this purpose, we utilized an NVIDIA A40 GPU, which provided the necessary computational power and memory. Additionally, to facilitate efficient processing of this large proteomic dataset, it was stored in the Zarr format codified by the Open Geospatial Consortium (https://www.ogc.org/).

#### HieVi Phage representation

To generate a vector representation of an entire phage proteome, we first ran pre-trained ESM-2 for each protein within each phage proteome and extracted the embeddings corresponding to each amino-acid in the protein sequence. Mean pooling over the amino-acid sequence was then used as our single protein representation. All proteins within a proteome were then normalized into unit vectors. We finally defined a Mean Phage Representation (MPR) by simply averaging over all ESM-2 protein representations within each proteome. The MPRs are thus vector representations of phage’s proteomes within the same 2,560-dimensional space as the protein embedding representation from ESM-2. We generated MPRs for all unique accession numbers in the HieVi dataset, resulting in 24,362 vectors.

#### HieVi Distances and Metrics

Euclidean distances between phage representations were used unless specified. Silhouette score is used to quantify the clustering validity of phage representations with respect to ICTV genus and subfamily annotations within annotated ICTV families^20^. For external validation with respect to the ICTV genus/subfamily annotations or vConTACT v2.0 VC clusters, we used Adjusted Mutual Information (AMI)^21^.

#### HieVi Clustering phage representations

For clustering the 24,362 MPRs, we used HDBSCAN (Hierarchical Density-Based Spatial Clustering of Applications with Noise), a clustering algorithm that builds on the DBSCAN (Density-Based Spatial Clustering of Applications with Noise) framework^22,23^. This allows for MPRs clustering based on density with a possibility of extracting flat clusters for a distance threshold. This distance threshold for extracting flat DBSCAN clusters is denoted as eps. Visualization of the high-dimensional phage representation in a two-dimension Phage Atlas was performed using Uniform Manifold Approximation and Projection (UMAP)^24^.

#### Phage networks visualization

For vConTACT v2.0 Phage similarity network and HieVi hierarchical tree visualization, we used Cytoscape 3.10.3^25^ completed with the free release of yFiles Hierarchical Layout algorithm from yWorks GmbH. The annotation file provided with the INPHARED dataset comprises for each virus accession number numerous information such as the current viral taxonomy, isolation host, genome length or number of tRNA genes. We enriched this table, whenever possible, with partial host taxonomy, host envelope descriptor (Gram positive, Gram negative or other descriptors) or virion morphology. We also extracted from protein annotations several keywords such as integrase, excisionase, RNA polymerase, etc. We added in this table for each accession number vConTACT v2.0 results as well as our own categories discussed in several Figures in this manuscript. This HieVi annotation table is used in Cytoscape to search and investigate HieVi hierarchical tree topology on various criteria and provided as a Supplementary File.

## Results

### HieVi Phage Atlas: phage clusters align with annotated genus

The 24,362 MPRs vectors representing all viruses in the HieVi dataset in the 2,560 dimensions space were projected on a two-dimensional HieVi Phage Atlas to visualize the viral diversity using UMAP (Figure 1).

**Figure 1:**
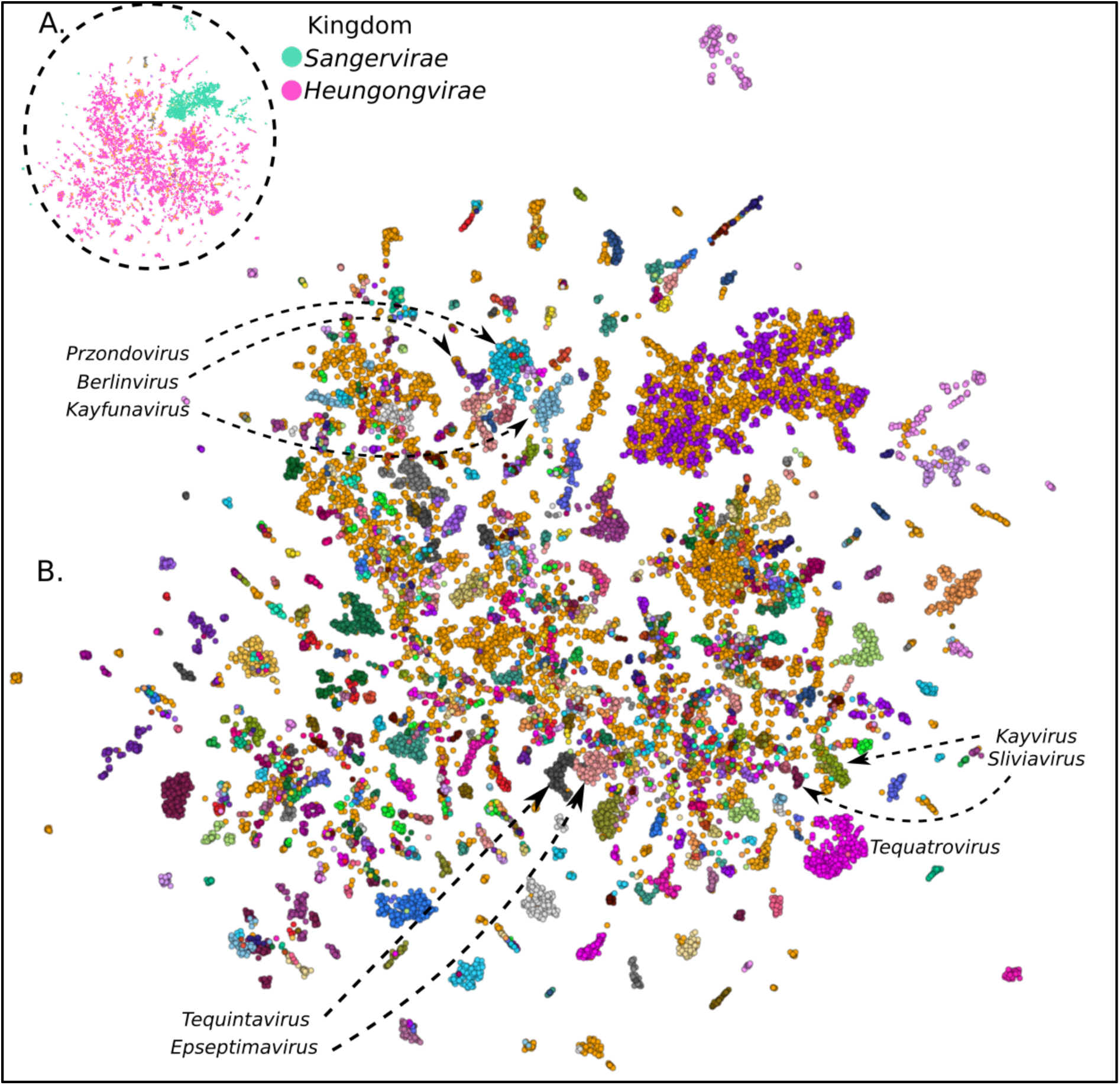
HieVi Phage Atlas of the HieVi dataset. Visualization of the phage representations (*n* = 24,362 viruses) in two dimensions. (A) Viruses colored by ICTV kingdoms. *Heunggongvirae* and *Sangervirae* ICTV kingdoms were highlighted in pink and green, respectively. Unclassified viruses are colored in orange. (B) Viruses colored by ICTV genera. Pointed clusters: *Prondzovirus*, *Berlinvirus* and *Kayfunavirus* (*Autographiviridae* family, *Studirevirinae* subfamily) ; *Tequintavirus* and *Epseptimavirus* (*Demerecviridae* family, *Markadamsvirinae* subfamily) ; *Silviavirus* and *Kayvirus* (*Herelleviridae* family, *Twortvirinae* subfamily) ; *Tequatrovirus* (*Straboviridae* family, *Tevenvirinae* subfamily). Unclassified viruses are colored in orange. An interactive HieVi Phage Atlas (September 2024 release) is available (see Data availability section).

Each point in this atlas is a phage genome from the HieVi dataset colored by an ICTV taxonomic rank. In Figure 1A, phages are colored by ICTV kingdoms. We observe that viruses are well clustered according to the realm they belong to, with *Heunggongvirae* and *Sangervirae* forming the two major clusters in the Phage Atlas. *Heunggongvirae* (dsDNA phages coding for a HK97-fold capsid protein) is the only kingdom from the *Duplodnaviria* realm (dsDNA phages) and encompasses the totality of the *Caudoviricetes* class (tailed phages), the most abundant group of phages in our dataset. *Sangervirae* (single-stranded DNA or ssDNA prokaryotic viruses coding for a single jelly-roll fold capsid protein) is one of the four kingdoms of the *Monodnaviria* realm (ssDNA phages). In Figure 1B, viruses are colored by ICTV genera. On the finer grain, well separated clusters can be observed that are consistent with the ICTV genus level classification in most cases. For example, *Tequatrovirus* forms a well separated cluster. *Epsetimavirus* and *Tequintavirus,* the two genera defining the *Markadamsvirinae* subfamily, form two separate clusters but remain close in the approximated manifold of MPRs. The clustering efficiency of the MPRs with respect to ICTV genus is quantified by a Silhouette score^20^ of 0.30. Silhouette score ranges from -1 to 1 and a score greater than zero indicates a fairly good clustering of MPRs at the genus level. It is worth keeping in mind that the unclassified viruses are ignored and that we trust the genus annotation.

HieVi Phage Atlas suggests that MPRs embedded multiscale information for classification of phages well-above the genus level. The ICTV genus is well defined by a single criterion, *i.e.* > 70% Average Nucleotide Identity (ANI) between genomes. With such a threshold, phage proteomes are thus expected to share a significant number of protein orthologs within a given genus. It is thus unsurprising that the MPRs cluster together by genus as these phage representations contain information about accumulated protein functions within a proteome. In analogy with natural language models, pre-trained mean sentence embeddings contain average semantic meaning of short documents and have been used as weak document classifiers. It is less obvious for subfamilies as they are not defined by strict criteria such as ANI but emerge from a combination of approaches such as gene/protein sharing networks or single/multiple gene/protein phylogenies.

To further understand why the simple averaging of all protein representations within a phage proteome led to well-defined phage clusters in the HieVi Phage Atlas, we drew in Figure 2 conceptual parallels with the methodology employed by vConTACT v2.0, a bioinformatic pipeline that cluster viruses according to their shared protein content^16^.

**Figure 2:**
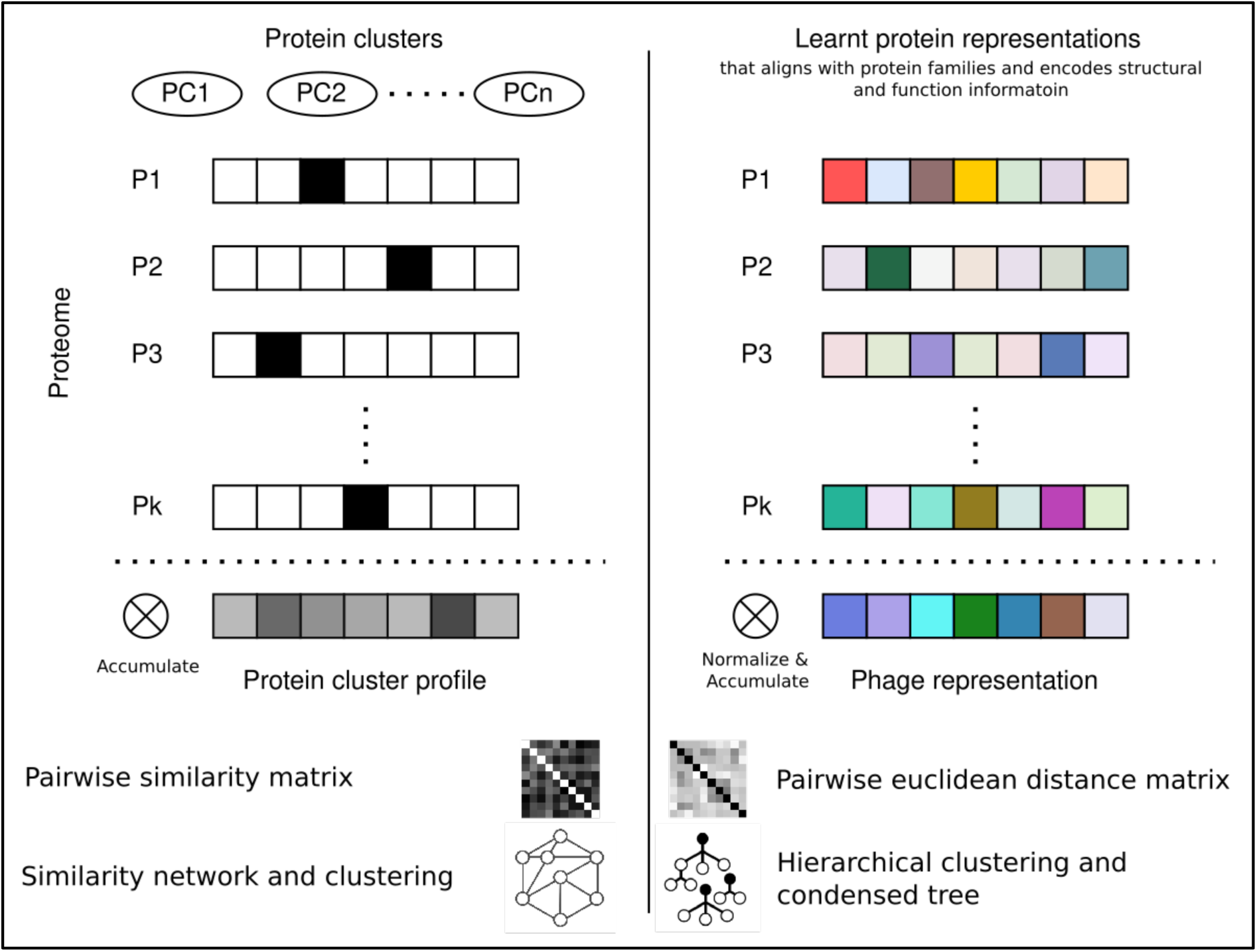
Conceptual parallels between vConTACT v2.0 and HieVi. On the one hand, in vConTACT v2.0, each protein may be seen as a one-hot vector in the protein cluster space with dimension (25,513 in the initial vConTACT v2.0 paper^16^). Then, the protein cluster profile for each genome is generated by summing over all the vectors. A similarity matrix is then computed using the hypergeometric formula (taking into account the genome lengths) and then a dense similarity network is generated and clustered using ClusterOne. On the other hand, in HieVi the protein representation generated using a pre-trained pLM has been shown to align with protein families and encodes protein function, structure and evolutionary fitness. Averaging over the protein representations in a proteome then gives a unique phage representation. A pairwise Euclidean distance matrix between all viruses is computed, then hierarchically clustered using HDBSCAN and a tree-like hierarchical network is finally generated.

As mentioned previously, vConTACT v2.0 pipeline first generates protein clusters (PCs) with MCL. Then for each proteome, protein cluster profiles were defined by accumulating the shared protein clusters. This is equivalent to representing proteins as one-hot vectors in the protein cluster space and then accumulating the proteins within a proteome to obtain a protein cluster profile which is a vector of size of the number of protein clusters (125,242 in this study with the HieVi dataset). A pairwise similarity matrix was then computed using these vectors and a similarity network was generated. Network clustering by ClusterOne was finally used to identify well separated clusters. Noteworthy, the number of protein clusters (PCs) increases in vConTACT v2.0 together with the number of genomes (2,304 genomes and 25,513 PCs in the initial vConTACT v2.0 paper^16^, 24,362 genomes and 125,242 PCs in the present study).

In HieVi, identification of a finite number of protein clusters isn’t necessary since the embeddings already encode protein family information in the entire space of the proteins in the trained dataset. Thus, adding new proteomes in HieVi is easy since it does not require creating new protein clusters. The protein representations are accumulated by simple averaging to generate MPRs. A pairwise Euclidean distance can then be generated for the entire dataset to build a network and clusters. However, we chose to employ hierarchical clustering and obtained a hierarchical network retaining multiscale relationships between phages in the dataset. In the next sections, we describe HieVi hierarchical clustering and the insights that it brings for phage classification.

#### HieVi hierarchical clustering and tree

We used HDBSCAN with the Euclidean distance metric between vectors to obtain a granular cluster so that there are at least two leaves per cluster, essentially constructing a dendrogram. HDBSCAN is designed to handle data with varying cluster densities and is particularly useful for identifying clusters of varying shapes and sizes without knowing the number of clusters apriori^23^. We then extracted the condensed tree that HDBSCAN uses to cluster the 24,362 viruses as our hierarchical tree of phages (Figure 3).

**Figure 3:**
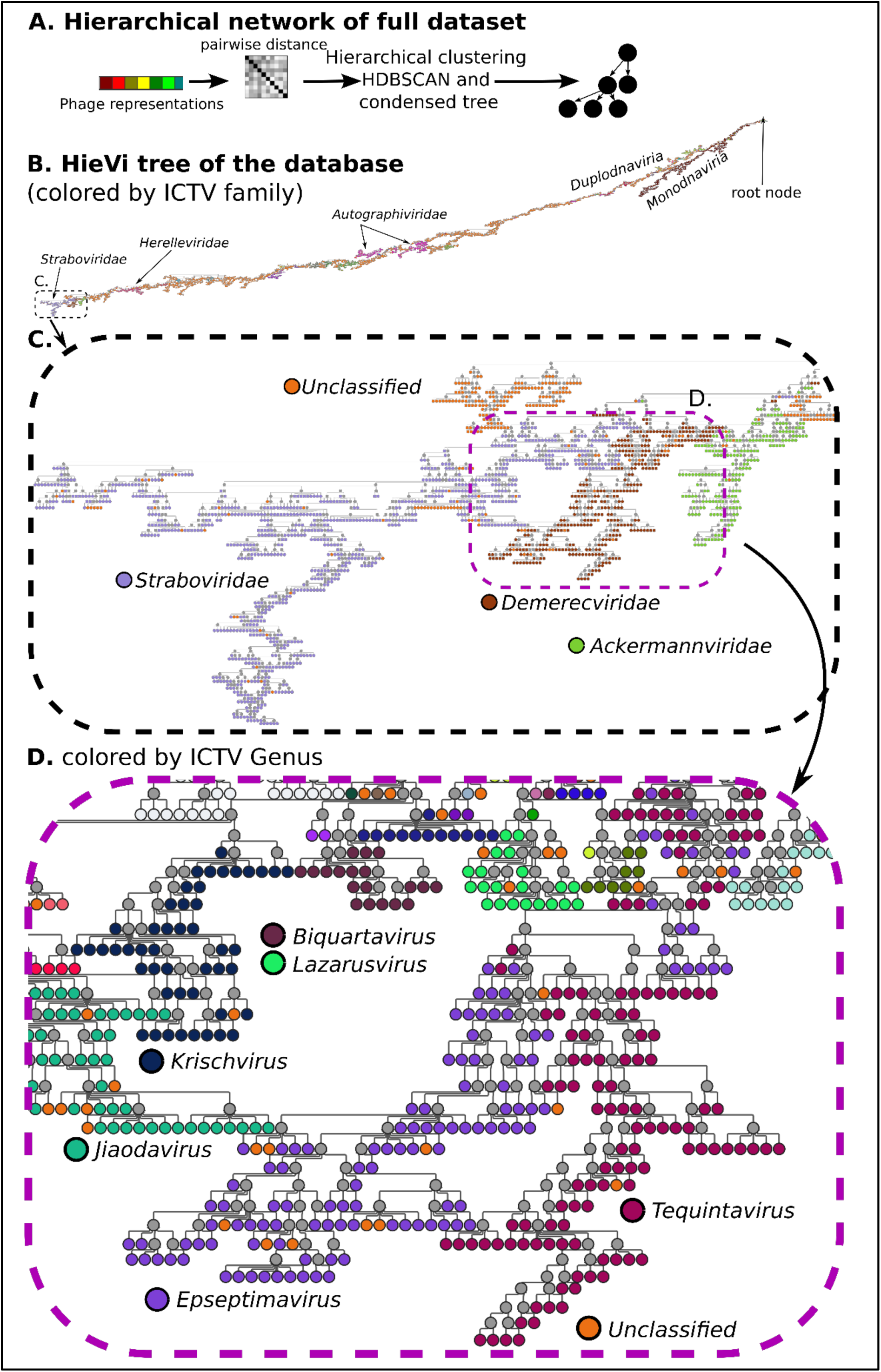
HieVi hierarchical network. (A) Workflow to generate the HieVi hierarchical tree from the MPRs. Network hierarchy and topology show multiscale phage groupings, coherent with the ICTV realm (*Duplodnaviria* and *Monodnaviria* in B), family (*Demerecviridae*, *Straboviridae* and *Ackermannviridae* in C) and genus (*Biquartavirus*, *Lazaarusvirus*, *Kirschvirus*, *Jiaodavirus*, *Epseptimavirus* and *Taquintavirus* in D). Relevant annotations are summarized in the Supplementary Excel file HieVi_annotations.xlsx. The HieVi hierarchical tree network can be downloaded and view in Cytoscape (see Data availability section)

Figure 3B shows an overview of the large network containing all virus proteomes in the dataset as a hierarchical tree. At the top of the hierarchy one can find the *Monodnaviria* (ssDNA phages) realm that transitions into phages from other realms (not indicated) and finally into *Duplodnaviria* (dsDNA phages). The large network is colored by ICTV families with orange representing unclassified viruses. For aid, we indicated *Autographiviridae*, *Herelleviridae* and *Staboviridae* families that we chose to study in further detail. On a finer scale in Figure 3C, we observe that phages are grouped in separated branches corresponding to the ICTV family. Finally at the finest grain, Figure 3D shows that phages cluster with respect to the ICTV genus. At least qualitatively, we see multiscale phage grouping emerging HieVi hierarchy and topology, which constitute a clarifying step beyond the vConTACT v2.0 Phage network for the entire dataset (Figure S1).

To further investigate the meaning of this topology, we first focused on the *Straboviridae* and *Herelleviridae* families. These two families were defined quite recently using whole genome approaches (see ICTV proposal 2021.082B for *Straboviridae* and Barylski et *al.* for *Herelleviridae*^13^). *Straboviridae* phages infect mostly *Gammaproteobacteria* belonging to *Enterobacterales*, *Aeromonadales*, *Moraxellales* and *Vibrionales* orders whereas *Herelleviridae* infect bacteria mostly belonging to *Bacillales* and *Lactobacillales* orders. Both families form several well-defined viral clusters in close proximity in the vConTACT v2.0 Phage Network (Figure S1).

For each of these two families, we identified in the HieVi hierarchical tree the top node of phages annotated for the corresponding ICTV family and extracted all its successors. We thus obtained one branch regrouping 94.8% of the *Straboviridae* phages and one branch regrouping 94.2% of the *Herelleviridae* phages. These two branches are described in Table 3.

**Table 3:** *Straboviridae* and *Herelleviridae* branches annotations. *Straboviridae* branch comprises 37 ICTV genera (64 phages are unclassified at the genus level). *Herelleviridae* branch comprises 36 ICTV genera (97 phages are unclassified at the genus level).

| Branch | ICTV /<br>Unclassified | Subfamily | Main host orders |
| --- | --- | --- | --- |
| *Straboviridae* (946) | 878 / 68 | *Guernseyvirinae* (1), *Tevenvirinae* (615), *Twarogvirinae* (34), Unclassified (296) | *Enterobacterales* (834), *Moraxellales* (42), *Aeromonadales* (40), *Vibrionales* (26) |
| *Herelleviridae* (517) | 528 / 43 | *Andregratiavirinae* (2), *Bastillevirinae* (149), *Brockvirinae* (51), *Jasinkavirinae* (36), *Joanriponvirinae* (9), | *Bacillales* (517), *Lactobacillales* (53), |
|  |  | *Sejongvirinae* (7), *Spounavirinae* (27),<br>*Twortvirinae* (264), Unclassified (26) |  |

Figure 4 displays the *Straboviridae* and *Herelleviridae* HieVi branches. We extracted as well the corresponding vConTACT v2.0 networks. Phages in both HieVi branches and vConTACT v2.0 networks were colored according to the ICTV genus.

**Figure 4:**
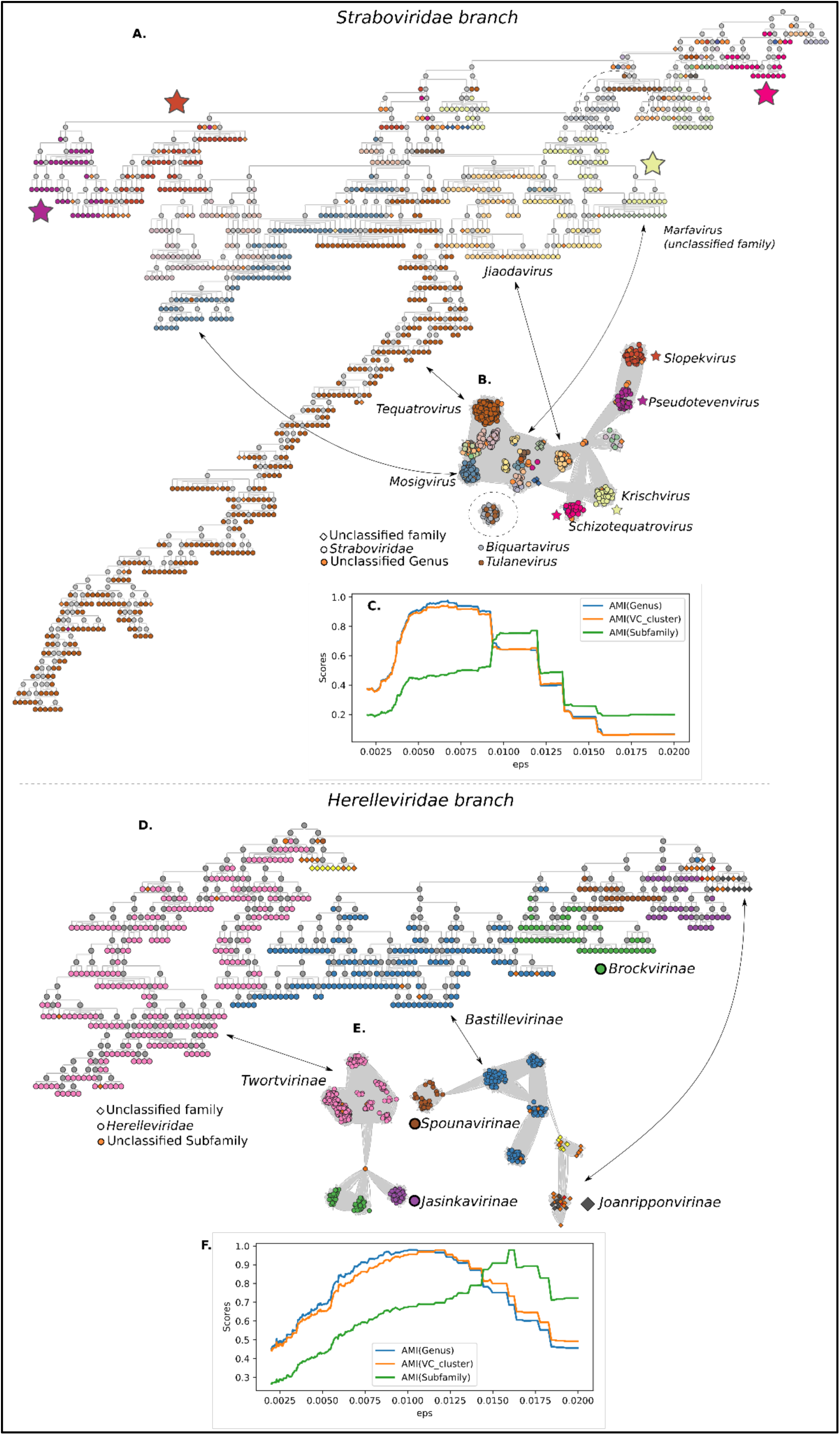
*Straboviridae* (left) and *Herelleviridae* (right) family. (A) HieVi branch for *Straboviridae* phages, colored by ICTV genera. (B) vConTACT v2.0 network for *Straboviridae* phages, colored by ICTV genera. (D) HieVi branch for *Herelleviridae* phages, colored by ICTV subfamilies. (E) vConTACT v2.0 network for *Herelleviridae* phages, colored by ICTV subfamilies. Some genus or subfamily correspondence between HieVi branches and vConTACT v2.0 networks are indicated by arrows. (C, F) Adjusted Mutual Information (AMI) scores plotted against eps distances with respect to ICTV genus (blue curves) and ICTV subfamily (green curves) for HieVi branches, with respect to Viral Cluster (VC) annotation (orange curve) for vConTACT v2.0 networks. (C) AMI scores for *Straboviridae* phages. (F) AMI scores for *Herelleviridae* phages. Relevant annotations are summarized in the Supplementary Excel file HieVi_annotations.xlsx.

Figure 4 shows that phages are well clustered in the HieVi hierarchical tree according to the ICTV genera (*Straboviridae*, Figure 4A) and the ICTV subfamilies (*Herelleviridae*, Figure 4D). This observation is quantitatively supported by the clustering efficiency metrics displayed in Figure 4C (*Straboviridae*) and Figure 4F (*Herelleviridae*). The AMI scores with respect to the ICTV genera (blue curves) and subfamilies (green curves) are > 0.8, indicative of an efficient clustering with respect to the annotations for these two families (AMI score varies from 0 to 1). We obtained the same performance for other phages families (Figure S2A). The AMI score profiles with respect to vConTACT v2.0 VCs (orange curves) are highly similar to the ones obtained with HieVi with the ICTV genus annotation, which is expected as it has been shown that VCs generally cluster well phages at the genus level. At variance with HieVi, vConTACT v2.0 VCs do not cluster phages at the subfamily level. For instance, *Bastillevirinae* subfamily phages are grouped in the *Herelleviridae* HieVi branch (highlighted in blue in Figure 4D) while they are separated in four VCs (VC_196, VC_197, VC_199 and VC_953, highlighted in blue in Figure 4E) in the vConTACT V2.0 network.

AMI score profiles displayed Figure 4C and 4E show that one can modulate the eps distance threshold to cluster phages at the genus or the subfamily level within a given family. Nevertheless, these AMI profiles also show that the eps value is not the same for a given taxonomic rank across all phage families. It is thus not possible to define a single eps threshold to predict genera (or subfamilies) across the entire HieVi hierarchical tree (Figure S2B).

As an intermediary conclusion, we evidenced that HieVi enables a representation of the phage world as we know it as a hierarchical tree that can capture multiscale relationships. On the broader view, viruses are rather well clustered according to the realm they belong to. We observed that viruses belonging to the same ICTV family remain generally close in the tree topology. We also qualitatively and quantitatively evidenced that HieVi is efficient in clustering viruses at two successive taxonomic ranks, *i.e*., genus and subfamily.

### Case study: *Autographiviridae*

According to ICTV, *Autographiviridae* is a phage taxon regrouping podovirus phages encoding their own single large subunit RNA polymerase with all genes transcribed on the Watson strand. Inclusion within this ICTV family is mostly defined by these two criteria (see ICTV proposal 2008.020-023B). *Autographiviridae* is a particularly illustrative example of the challenges in phage classification as well as how the HieVi tool could be useful in this context to biologists in the field for *Autographiviridae* phages taxonomy.

Our dataset comprises 1,743 *Autographiviridae* viruses assigned to eight subfamilies and 130 genera in the current ICTV classification, evidencing that this family is very diverse. In vConTACT v2.0 Phage network, *Autographiviridae* phages form several, unconnected clusters (Figure S1, highlighted in blue). At variance with the phage families we previously investigated, Autographiviridae phages are not clustered in the same branch in the HieVi hierarchical tree. 96.7% of these phages are spread among nine branches (01 to 09, Table 4), with 01 and 02 branches accounting for 83.6% of the *Autographiviridae* dataset.

**Table 4:** *Autographiviridae* branches annotations. These nine branches comprise in total 121 ICTV genera (382 phages are unclassified at the genus level). *Phages belong to another ICTV family.

| Branch | *Autographiviridae* / Unclassified / Other* | Subfamily | Main host orders | Predicted integrase Yes / No | Genome size (kb) |
| --- | --- | --- | --- | --- | --- |
| 01 (952) | 858 / 94 | *Studiervirinae* (843),<br>Unclassified (110) | *Enterobacterales* (827),<br>*Pseudomonadales* (68),<br>*Vibrionales* (25) | 1 / 951 | 39.7 ± 1.2 |
| 02 (671) | 600 / 70 / 1 | *Bclavirinae* * (1),<br>*Colwellvirinae* (19),<br>*Corkvirinae* (36),<br>*Krylovirinae* (75),<br>*Melnykvirinae* (36),<br>*Molineuxvirinae* (108),<br>*Okabevirinae* (17),<br>*Slopekivirinae* (127),<br>*Studiervirinae* (2),<br>Unclassified (250) | *Enterobacterales* (392),<br>*Pseudomonadales* (84),<br>*Xanthomonadales* (50),<br>*Vibrionales* (45),<br>*Burkholderiales* (34),<br>*Aeromonadales* (27),<br>*Hyphomicrobiales* (23),<br>*Caulobacterales* (11) | 22 / 649 | 43.6 ± 1.9 |
| 03 (121) | 101 / 20 / 0 | *Beijerinckvirinae* (91),<br>*Slopekivirinae* (7),<br>Unclassified (23) | *Acinetobacter* (107) | 0 / 121 | 41.5 ± 0.9 |
| 04 (50) | 50 / 0 / 0 | Unclassified (50) | *Prochlorococcus* (29),<br>*Synechococcus* (18) | 15 / 35 | 42.8 ± 4.0 |
| 05 (36) | 36 / 0 / 0 | Unclassified (36) | *Pelagibacter* (30) | 31 / 5 | 40.6 ± 2.0 |
| 06 (14) | 14 / 0 / 0 | Unclassified (14) | *Enterobacterales* (14) | 0 / 14 | 42.0 ± 0.7 |
| 07 (14) | 14 / 0 / 0 | Unclassified (14) | *Rhizobium* (4),<br>*Ralstonia* (8) | 14 / 0 | 41.7 ± 1.5 |
| 08 (9) | 9 / 0 / 0 | Unclassified (9) | *Roseobacter* (9) | 9 / 0 | 40.0 ± 0.9 |
| 09 (4) | 4 / 1 / 1 | *Studiervirinae* (4),<br>*Tunavirinae* * (1), | *Enterobacterales* (4) | 0 / 9 | 48.7 ± 11.8 |
|  |  | Unclassified (1) |  |  |  |

The corresponding HieVi branches and vConTACT v2.0 networks are displayed in Figure 5 with viruses colored by ICTV subfamilies.

**Figure 5:**
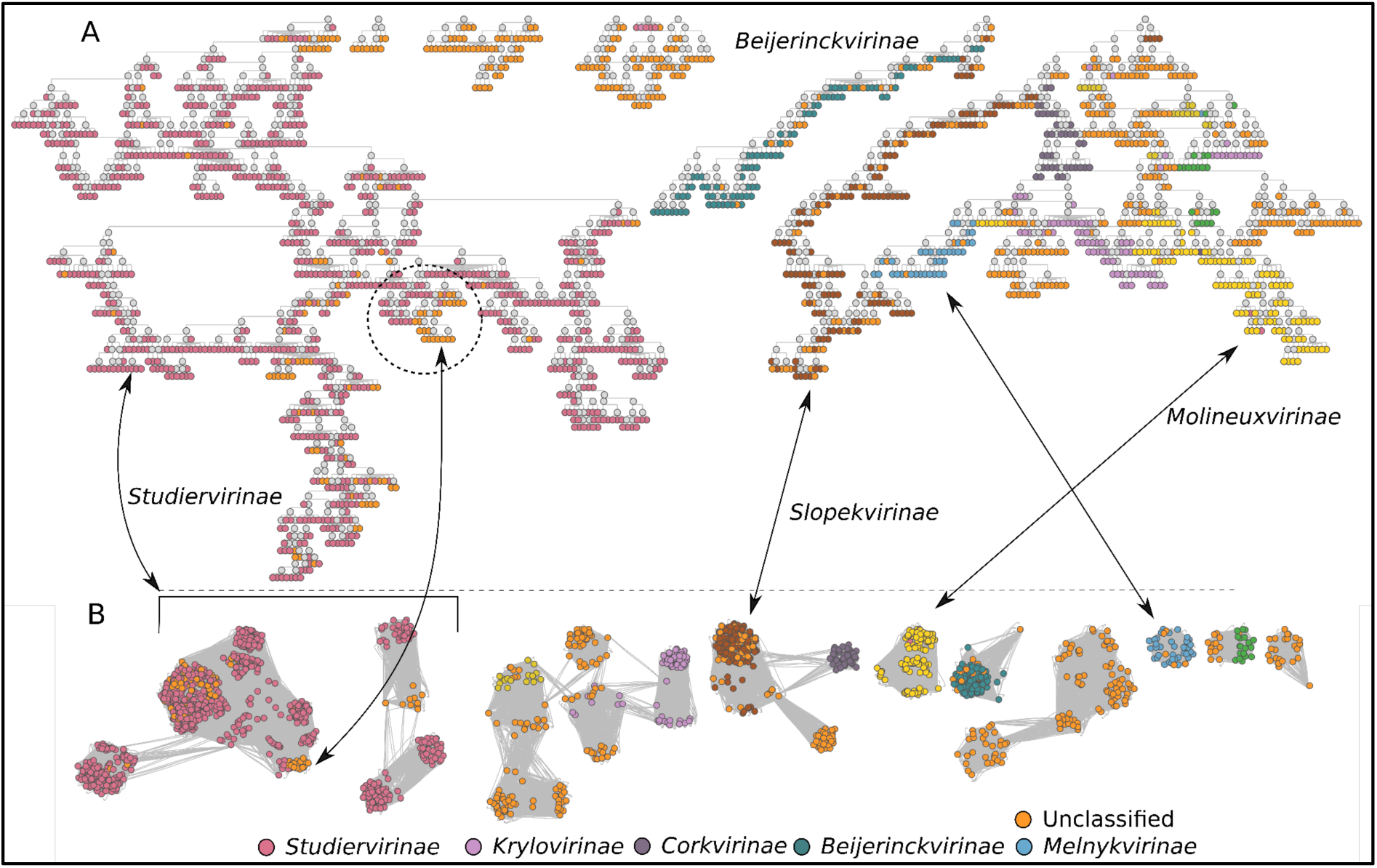
*Autographiviridae* family. A) HieVi branches for *Autographiviridae*. B) vConTACT v2.0 network for *Autographiviridae*. Phages are colored by ICTV subfamilies. Relevant annotations are summarized in the Supplementary Excel file HieVi_annotations.xlsx.

As was the case for *Straboviridae* and *Herelleviridae*, we observed that *Autographiviridae* phages cluster in the HieVi hierarchical tree according to the ICTV genus (data not shown) but also with the ICTV subfamily (Figure 5A). Such is not always the case in vConTACT v2.0 phage network. As an example, 843 out of the 856 *Studiervirinae* phages in the dataset are grouped into a single HieVi branch (Figure 5A) while they are spread among several poorly connected or unconnected viral clusters in vConTACT v2.0 network (Figure 5B). HieVi topology suggests new relationships between *Autographiviridae* subfamilies and HieVi branches could be used by experts in the field as a blueprint to investigate new taxa regrouping previously acknowledged ones. HieVi is thus able to capture connections between subfamilies that were previously unlinked.

On a finer grain, HieVi can help classify unclassified phages in an effective fashion as illustrated in Figure 6 within *Autographiviridae* branch 01 that mostly includes *Studiervirinae* phages.

**Figure 6:**
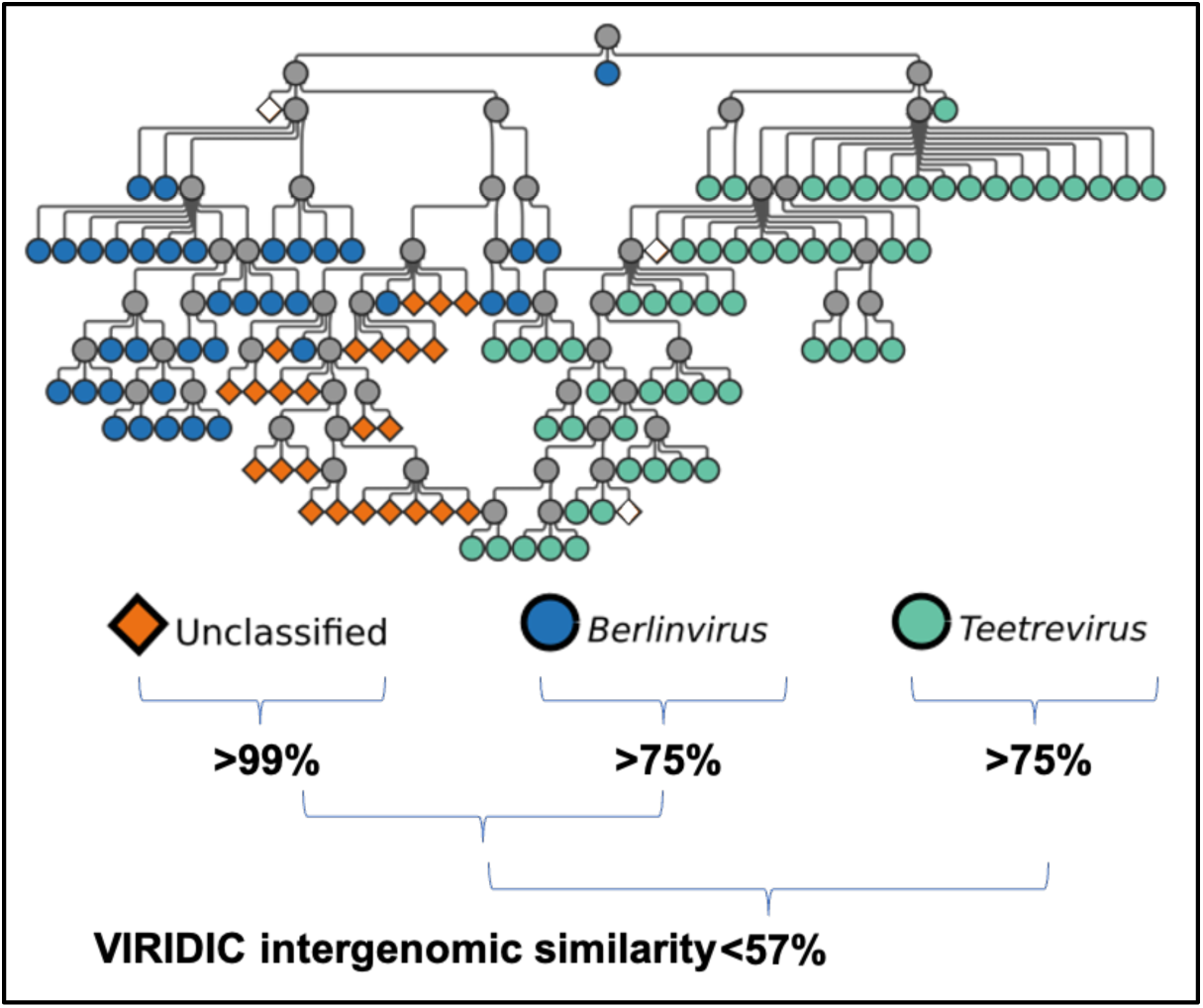
Taxonomic assignment of unclassified phages within the *Autographiviridae* branch 01 (*Studiervirinae*). (◆) Unclassified family, subfamily and genus. (⦁) *Berlinvirus*. (⦁) *Tetreevirus*. Intergenomic similarity between groups was computed with VIRIDIC. Relevant annotations are summarized in the Supplementary Excel file HieVi_annotations.xlsx.

In this Figure, one can identify a branch comprising 24 phages represented in orange, unclassified at the family, subfamily and genus levels, plugged into the *Berlinvirus* genus branch and adjacent to the *Treetrevirus* branch. Since we previously evidenced that HieVi topology follows genus and subfamily annotations, we posit these unclassified phages are, at the least, *Autographiviridae* belonging to the *Studiervirinae* subfamily, and eventually *Berlinvirus*. We computed with VIRIDIC^26^ the intergenomic distances matrix for these 24 unclassified phages and the surrounding *Berlinvirus* and *Tetreevirus* and clustered them with ANI threshold > 95% and >70% for species and genus levels, respectively. VIRIDIC clustering clearly states that these 24 unclassified phages represent a single, *Berlinvirus* species.

As mentioned previously, *Autographiviridae* is a diverse family with eight subfamilies and 130 genera. Combining HieVi topology, functional annotation or other differentiating criteria can also draw attention to new relationships. A first type of connection relates to *Autographiviridae* genome size. The second type derives from the combination of HieVi topology and protein functional annotation. For the latter we examined the presence or absence of a predicted integrase within the proteomes. Table 4 summarizes genome length and predicted integrase for all nine HieVi *Autographiviridae* branches.

*Autographiviridae* genome size distribution is bimodal, with one peak around 40.5 kb and the other around 43.5 kb (Figure S3). In Table 4 we can see that *Autographiviridae* phages are well sorted across the two main branches (01 and 02) according to their genome size. Branch 01 hosts smaller genomes (39.7±1.2 kb) than branch 02 (43.6±1.9 kb). are mostly clustered in two branches (01 and 03), while bigger genomes (>42 kb) are mostly clustered in branch 02. Genome size could thus be used as an additional criterion to improve *Autographiviridae* taxonomy.

Regarding the presence of a predicted integrase protein in *Autographiviridae* proteomes, we can readily observe in Table 4 that three branches (05, 07 and 09) almost exclusively encompass phages harboring an integrase, an indication of a putative temperate lifestyle. Branch 04 presents a mixed situation with about half of its members coding for an integrase, either due to simple functional misannotation or maybe, more interestingly, due to gain/loss of function.

On a broader perspective, our results suggest that HieVi tree topology can assist experts in the field in refining, improving or even questioning the current structure of the *Autographiviridae* family. Since HieVi branches regrouping subfamilies are consistent with established phylogenies, these branches could serve to define new families encompassing phylogenetically related phages coding for their own large single subunit RNA polymerase. Genome size or the presence of an integrase could also be used as criteria to define membership in these new families. *Autographiviridae* could thus be promoted at the rank order (*Autographivirales*) and include these newly defined families. The inclusion criteria would be the same as those defined for the current *Autographiviridae* family.

#### Case study: Discovery of a cohesive group of lambdoid phages

We identified in the HieVi hierarchical tree an interesting branch clustering 470 phages, 348 of which code for a predicted integrase necessary (but not sufficient) for a temperate lifestyle (Table 5). Kupcozk et *al.* recently investigated co-transferred genes in lambdoid phages and defined 12 groups. Within each of these 12 groups gene content is highly similar while it is highly divergent across the 12 groups^12^. Among these, one can find well studied temperate phages such as *Lambdavirus* Lambda, *Byrnievirus* HK97 and *Lederbergvirus* P22. The INPHARED dataset we used in our study comprises 30 lambdoid genomes out of the 31 used by Kupcozk et *al.* (not counting their *E. coli* prophage dataset that is not included in the HieVi dataset). Among these 30 lambdoid phages, 29 are included in this HieVi branch. We thus provisionally named this branch “Lambdoid branch” (Table 5). This branch also includes 186 temperate phages out of the 190 present in our dataset annotated as Shiga Toxin 1- and Shiga Toxin 2- converting phages.

**Table 5:** Lambdoid branch annotations. This branch comprises 32 ICTV genera (204 phages are unclassified at the genus level).

| Branch | Unclassified family | Subfamily | Main host order |
| --- | --- | --- | --- |
| 01 (470) | 470 | *Hendrixvirinae* (18), *Sepvirinae* (150),<br>Unclassified (302) | *Enterobacterales* (454) |

The HieVi lambdoid branch is presented in Figure 8 where phages are colored by ICTV genera.

**Figure 8:**
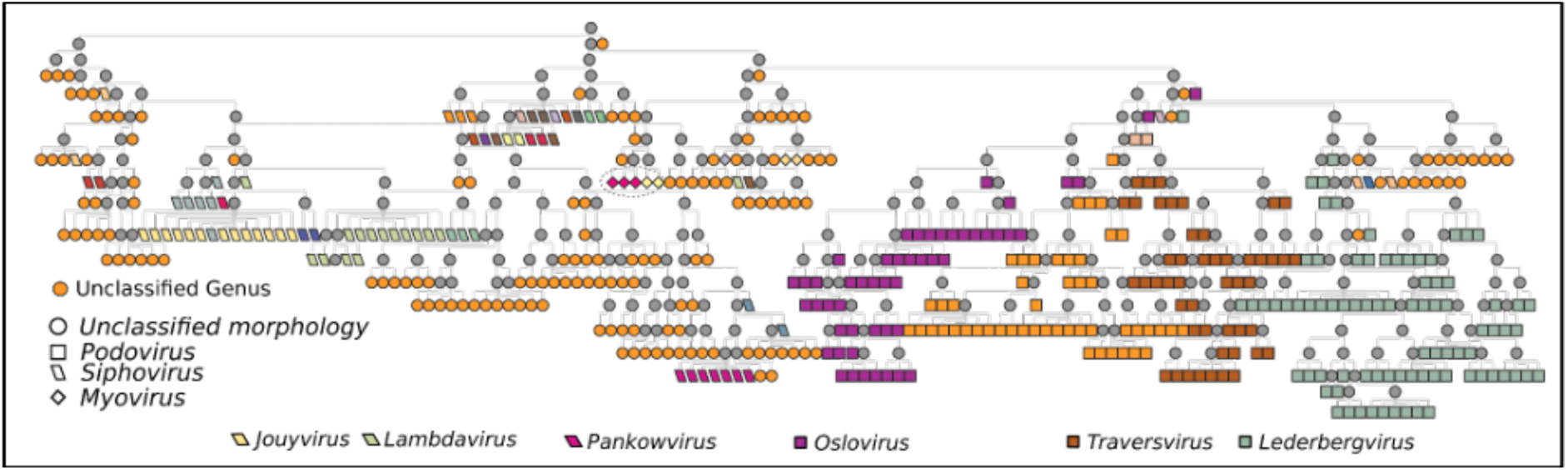
HieVi Lambdoid branch. Viruses are colored by ICTV genera. Virion morphology: (○) Unclassified, (□) Podovirus, (▱) Siphovirus, (◊) Myovirus. Highlighted genera: *Jouyvirus* (type phage Gifsy), *Lambdavirus* (type phage Lambda), *Pankowvirus*, *Oslovirus* (type phage TL2011), *Traversvirus* (type phage 933W) and *Lederbergvirus* (type phage P22). Relevant annotations are summarized in the Supplementary Excel file HieVi_annotations.xlsx.

All phages in this branch are unclassified at the family level and includes only two defined ICTV families, the *Sepivirinae* (including *Traversvirus* type phage 933W) and *Hendrixvirinae* (including *Byrnievirus* type phage HK97). As we observed previously in other HieVi branches, phages are well clustered according to the ICTV genus (Figure 8) and the two ICTV subfamilies (data not shown). Interestingly, this group contains different virion morphologies. For example, *Escherichia* phages HK97 and Lambda exhibit siphovirus morphology (long, flexible, non-contractile tail), *Salmonella* phage P22 and *Escherichia phage* HK620 are podovirus (short, non-contractile tail) and *Edwardsiella* phage GF-2 is a myovirus (long, contractile tail). In Figure 8 we can observe that phages in the Lambdoid branch cluster rather well according to their reported morphology: siphovirus (e.g., *Lambdavirus*, *Jouyvirus* and *Hendrixvirinae* phages) are mostly clustered at the left hand-side of the network while podovirus (*e.g.*, *Sepvirinae* and *Lederbergvirus* phages) are mostly clustered at the right hand-side.

Lambdoid phages have been well studied for decades and from these emerged the concept of mosaic genomes where genes are clustered in functional units that can be recombined between phages. As mentioned in the Introduction, this makes it difficult to trace the evolutionary trajectory of entire genomes. It is thus surprising that HieVi clustered all these lambdoid phages within a branch despite the gene content divergence across the 12 groups defined by Kupcozk et *al.* and with phages exhibiting various virion morphologies and a wide range of genome sizes (40 - 70 kb). We posit that HieVi can capture evolutionary relationships that extend beyond the genus and subfamily taxonomic ranks and probably offers the opportunity to identify recombinant groups.

### HieVi workflow for new genomes

HieVi pipeline is based on a precomputed Mean Phage Representation (MPRs) database, the HieVi database. This MPR database was generated from 24,362 annotated genomes in September 2024. We designed HieVi so that it is easily scalable (Figure 9). Placing few new genomes in the current HieVi tree hierarchy will simply require the generation of the corresponding MPRs through the pipeline. A search for the k-nearest neighbors will enable the user to quickly visualize these new genomes in the HieVi tree topology. For larger sets of data, the MPRs will be added to the HieVi database. The new Euclidean matrix will be computed and the MPRs clustered *de novo* with HDBSCAN to generate a new, updated HieVi hierarchical tree.

**Figure 9:**
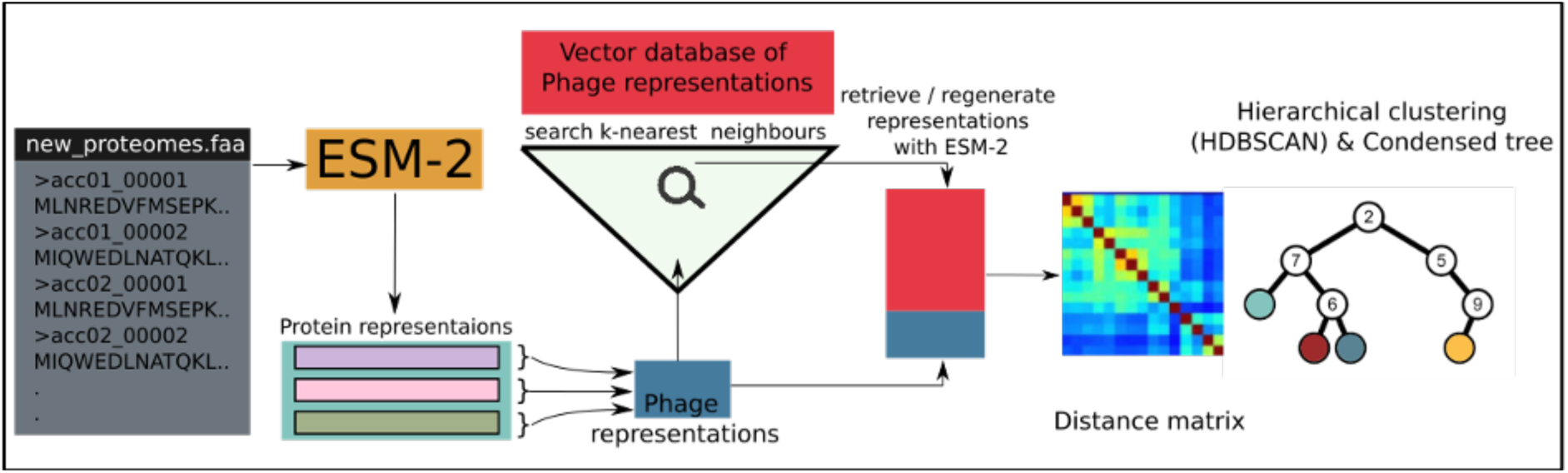
HieVi workflow for new proteomes. New viral proteins are first converted into vector representations with ESM-2 then normalized to unit length. Mean Phage Representations (MPR) vectors are then generated for each new virus by averaging all normalized protein vector representations within each new proteome. For small datasets we retrieve from our HieVi phage representation database the k-nearest neighbors. We then compute for this subset the Euclidean distance matrix, perform hierarchical clustering with HDBSCAN and finally generate the condensed tree.

As mentioned previously, every new MPR will be in the same dimensional space as the protein embedding space created by ESM-2. This makes HieVi easily scalable as there is no need to re-process the genomes already contained in the HieVi database, which is not the case for example with vConTACT v2.0. The processing of new genomes through the HieVi pipeline described in Figure 9 will result in an updated HieVi database containing the new dataset MPRs.

## Discussion

In this work, we explore the application of protein Language Models for the comparison and classification of phages, presenting a novel approach which extends the repertoire of proteome-based analysis of phages. We introduced *Hierarchical Viruses* or HieVi, a method that has the potential to address the increasing need for managing and organizing large datasets of phage genomes beyond multiple sequence alignment methods.

Leveraging pLMs and density based hierarchical clustering, HieVi produces a hierarchical tree rather than a dense network which facilitates easier visualization and interpretation of phage relationships in large datasets. HieVi also allows for new viromes to be classified and organized around known and well-studied complete genomes of viruses.

We showed that the HieVi tree organizes viruses into a hierarchical structure where multi-scale taxonomic ranks emerged, aligning well with current ICTV annotation (Figure 3). Through selected *Caudoviricetes* phage families - the *Herelleviridae*, *Straboviridae* and *Autographiviridae* - we first qualitatively observed that phages are grouped in the HieVi hierarchical tree topology along their ICTV genera and subfamilies (Figures 3 to 5 & 8). Clustering efficiency metrics quantitatively supported these observations (Figure 4 & S2). Phages belonging to the same ICTV family are also grouped in the tree topology (Figure 3), with the notable exception of *Autographiviridae*. Existing phylogeny within families has been defined using various approaches, the most recent one relying on whole genome/proteome analyses. Since the HieVi hierarchy is generally in good accordance with these established phylogenies, we posit that this hierarchy encodes phylogenetic relationships, at least up to the family level and can assist biologists to discover and define new phage families.

The *Autographiviridae* family, essentially defined by the presence of a gene encoding a large single subunit RNA polymerase, is an illustrative example of how the HieVi hierarchical tree topology can reveal unsuspected complexity within a taxon. Previous studies have evidenced the diversity of this family, with 130 genera and eight subfamilies. At variance with other phage families, the *Autographiviridae* does not appear as a single branch in the HieVi tree topology but is split into nine distinct branches. Within each branch, genus and subfamily clustering is respected as evidenced in other phage families, suggesting that each branch represents a phylogenetic unit on its own (Figure 5). Combining tree topology, functional annotation (*e.g.*, the presence of an integrase) and other genomic features (*e.g.*, genome size), the nine HieVi branches encompassing the *Autographiviridae* can serve as a blueprint for phage taxonomists to define nine new monophyletic taxa regrouped at a new, higher taxonomic rank, the *Autographivirales* order.

Latest phage phylogeny relies on the convergence of several whole genome/proteome analyses as exemplified by the *Herelleviridae* family^13^. Thus, we demonstrate that HieVi enables the emergence of a multi-scale taxonomic organization of phages, a feature that is not readily apparent in tools like vConTACT v2.0. Within ICTV families, phages share a significant number of proteins orthologs. The organization of the lambdoid branch we defined suggests that the HieVi hierarchical tree captures more than just shared genes (Figure 8). With this example, we showed that HieVi transcends conventional gene-sharing metrics and highlights evolutionary insights into the gene flux resulting in phage genome mosaicism.

We thus hypothesize that pLM-based phage comparison and classification enabled by HieVi enhances our ability to discover and interpret evolutionary patterns, possibly phylogeny-driven view of phage diversity. Large proteomic datasets, with ever-increasing additions due to global efforts in viral ecology, require better data organization. We posit that HieVi, a scalable and fast method for classification, equips researchers in the field of phage studies with a powerful tool for organizing, classifying, and exploring the complex and ever-growing landscape of viral ecology, paving the way for future discoveries.

Conceptually, this study builds on the advancements seen in protein Language Models and the ESM Metagenomic Atlas published only a year ago^5^. Protein Language Models have shown an impressive ability to learn the ’language’ of proteins in a self-supervised manner, providing insights into their functions and evolutionary relationships^27^. Since a genome code for a set of proteins, we hypothesized and showed that such models could be harnessed to learn not just protein functions but also how they assemble to define the proteomic capabilities of a biological entity as a whole. HieVi serves as an initial example of leveraging this approach to compare and classify biological entities. This advancement offers important perspectives in comparative genomics and related fields. As a perspective, HieVi could be refined to better address challenges associated with metagenomic sequences, including large datasets, partial sequences, and contamination by non-viral nucleic acids. Enhancements might involve fine-tuning the underlying protein language models specifically on viral proteins for diverse tasks. Additionally, since HieVi is built on the embedding space of viral proteins, it inherently captures functional and evolutionary information that can be further leveraged to explore new research directions. The adoption of larger and more advanced protein language models is also expected to improve HieVi accuracy and provide deeper insights into evolutionary relationships.

## Data availability

HieVi UMAP Phage Atlas September 2024 release (Figure 1) can be downloaded at https://github.com/pswapnesh/HieVi/raw/refs/heads/main/HieVi_UMAP.html then interactively viewed in a web browser. Annotation tables are available as a Supplementary Excel File (HieVi_annotations.xlsx). The annotated HieVi hierarchical tree is available as a Supplementary file in Cytoscape format (HieVi_Hierarchical_tree.zip).

Due to the large file size, the vConTACT v2.0 similarity network (Figure S1) is available on request. All vConTACT v2.0 results are included in the above-mentioned HieVi annotation file.

## Supporting information

HieVi annotation tables

Cytoscape HieVi Hierarchical tree file

## Acknowledgments

We are thankful to all present and past members of the “Phage Cycle and Bacterial Metabolism” group at Laboratoire de Chimie Bactérienne (LCB) for stimulating discussions and constant interest. We are grateful to Leon Espinosa (LBC) for fruitful discussions during the design of this project. The Centre National pour la Recherche Scientifique (CNRS), Aix-Marseille Université (AMU) and the Institut de Microbiologie de la Méditerranée (IMM) support our research. LCB and IMM provided computational resources.

## Author Contributions

SP, MA and NG designed the project. SP implemented machine learning approaches. NG performed comparative genomics. SP, MA and NG wrote the manuscript.

## Supplementary files

- HieVi_hierarchical_tree.zip: The annotated HieVi hierarchical tree of viruses file in Cytoscape format.
- HieVi_annotations.xlsx: This Excel file contains 1) the INPHARED annotation table (September 2024 release), 2) the HieVi hierarchical tree annotation table (September 2024 release), 3) the *Straboviridae* branch 01 annotation table, 4), the *Herelleviridae* branch 01 annotation table, 4) the *Autographiviridae* branches 01 to 09 annotation table, 5) *Berlinvirus* and *Tetreevirus* branch annotation table and 6) the Lambdoid branch 01 annotation table.

---

**Figure S1:**
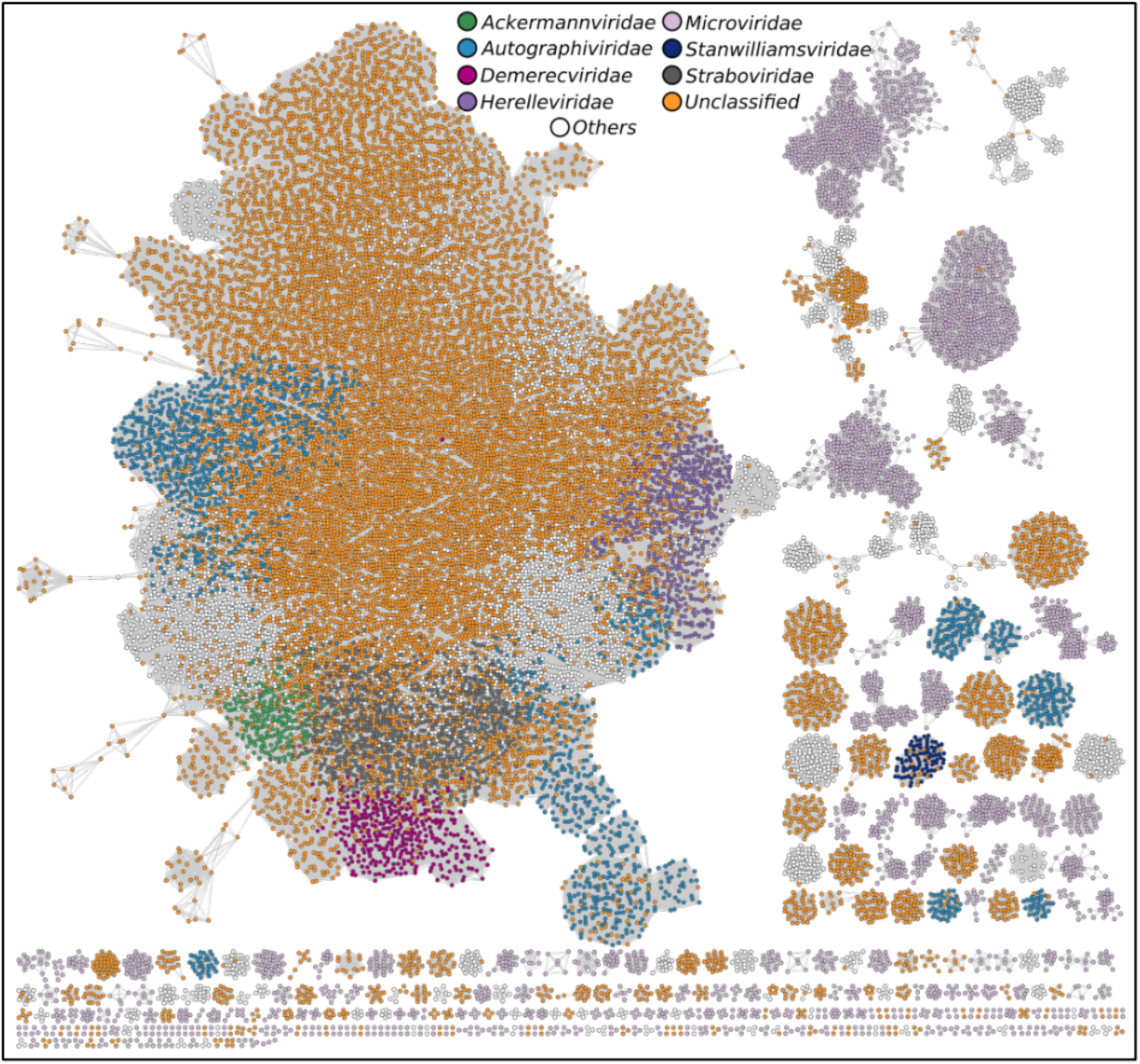
vConTACT v2.0 Phage Network. *Straboviridae*, *Herelleviridae* and *Autographiviridae* as well as several other ICTV families are colored (see legend). Phages belonging to other ICTV families are colored in white. Unclassified families are colored in orange. Relevant annotations are summarized in the Supplementary Excel file HieVi_annotations.xlsx.

**Figure S2:**
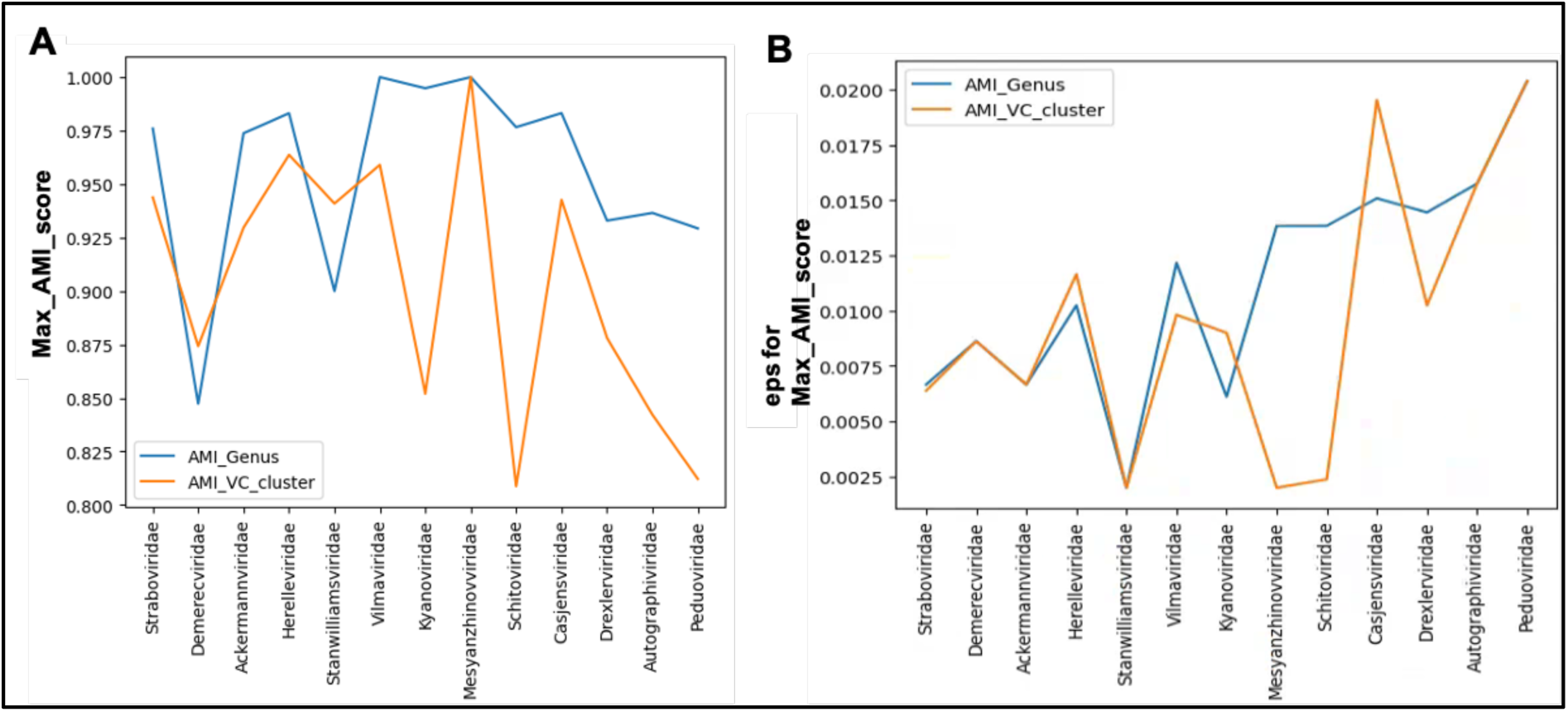
HieVi clustering efficiency across 13 ICTV families. A) AMI maximum score with respect to ICTV genus (blue line) and vConTACT v2.0 VCs (orange line). B) eps distance threshold for maximum AMI score with respect to ICTV genus (blue line) and vConTACT v2.0 VCs (orange line).

**Figure S3:**
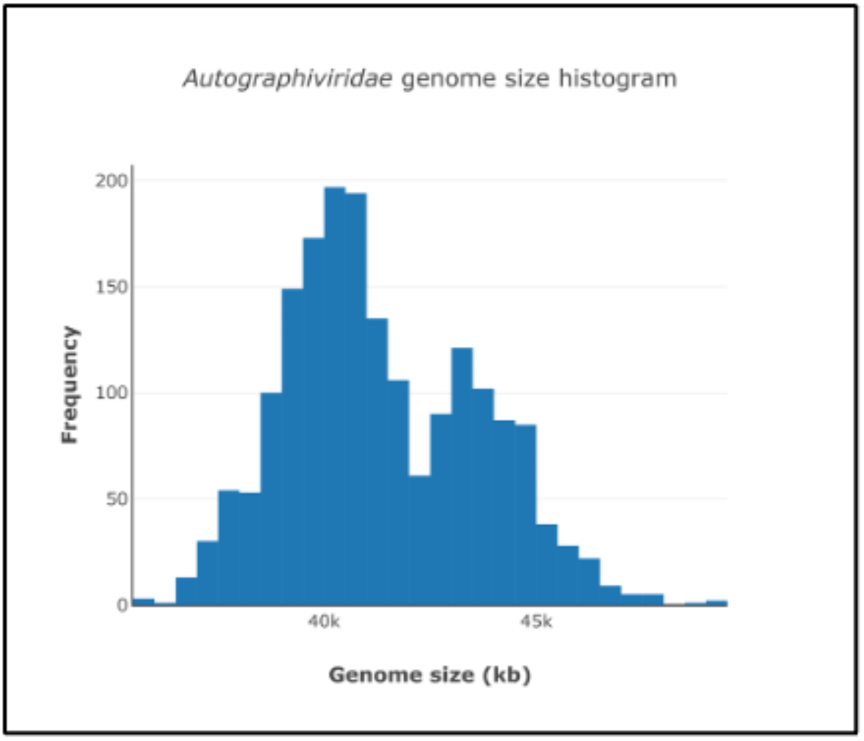
*Autographiviridae* genome size distribution. Histogram of *Autographiviridae* genome sizes included in branches 01 to 09. Bin size = 0.5 kb. Genome sizes are summarized in the Supplementary Excel file HieVi_annotations.xlsx.

